# PRDM6 promotes medulloblastoma by repressing chromatin accessibility and altering gene expression

**DOI:** 10.1101/2023.08.29.555389

**Authors:** Christin Schmidt, Sarah Cohen, Brian L. Gudenas, Sarah Husain, Annika Carlson, Samantha Westelman, Linyu Wang, Joanna J. Phillips, Paul A. Northcott, William A. Weiss, Bjoern Schwer

**Affiliations:** Department of Neurological Surgery, University of California San Francisco, San Francisco, CA, USA; Department of Cellular and Molecular Pharmacology, University of California, San Francisco, San Francisco, CA, USA; Eli and Edythe Broad Center of Regeneration Medicine and Stem Cell Research, University of California, San Francisco, San Francisco, CA, USA; Weill Institute for Neuroscience, University of California, San Francisco, San Francisco, CA, USA; Helen Diller Family Comprehensive Cancer Center, University of California, San Francisco, San Francisco, CA, USA; Bakar Aging Research Institute, University of California, San Francisco, San Francisco, CA, USA; Kavli Institute for Fundamental Neuroscience, University of California, San Francisco, San Francisco, CA, USA; Division of Brain Tumor Research, Department of Developmental Neurobiology, St. Jude Children’s Research Hospital, Memphis, TN, USA; Department of Neurology, University of California San Francisco, San Francisco, CA, USA; UCSF Tumor SPORE Biorepository, University of California, San Francisco, San Francisco, CA, USA

## Abstract

*SNCAIP* duplication may promote Group 4 medulloblastoma *via* induction of PRDM6, a poorly characterized member of the *PRDF1 and RIZ1 homology domain-containing* (PRDM) family of transcription factors. Here, we investigated the function of PRDM6 in human hindbrain neuroepithelial stem cells and tested PRDM6 as a driver of Group 4 medulloblastoma. We report that human PRDM6 localizes predominantly to the nucleus, where it causes widespread repression of chromatin accessibility and complex alterations of gene expression patterns. Genome-wide mapping of PRDM6 binding reveals that PRDM6 binds to chromatin regions marked by histone H3 lysine 27 trimethylation that are located within, or proximal to, genes. Moreover, we show that PRDM6 expression in neuroepithelial stem cells promotes medulloblastoma. Surprisingly, medulloblastomas derived from PRDM6-expressing neuroepithelial stem cells match human Group 3, but not Group 4, medulloblastoma. We conclude that PRDM6 expression has oncogenic potential but is insufficient to drive Group 4 medulloblastoma from neuroepithelial stem cells. We propose that both PRDM6 and additional factors, such as specific cell-of-origin features, are required for Group 4 medulloblastoma. Given the lack of PRDM6 expression in normal tissues and its oncogenic potential shown here, we suggest that PRDM6 inhibition may have therapeutic value in PRDM6-expressing medulloblastomas.

## INTRODUCTION

Medulloblastoma, a malignant childhood tumor of the cerebellum, represents a clinically and molecularly heterogeneous cancer originating from aberrant hindbrain development^1–4^. The cellular origins and genetic drivers of WNT and SHH medulloblastomas are relatively well characterized but drivers of Group 3 and 4 medulloblastoma remain unclear. Especially Group 4 medulloblastomas show a complex range of molecular aberrations and clinical outcomes^4,5^.

One of the most characteristic alterations of Group 4 medulloblastoma is the expression of PRDM6, which occurs in 17% of all Group 4 medulloblastoma cases^1,4^. Moreover, PRDM6-expressing Group 4 medulloblastomas often show other alterations, such as aberrant expression of the *MYCN* oncogene^5^. PRDM6 is a poorly characterized member of the *PRDF1 and RIZ1 homology domain-containing* (PRDM) family of transcription factors that possess histone lysine methyltransferase activities through their catalytic SET domain^6^. However, the enzymatic activity and molecular function of human PRDM6 is unclear. Based on findings in other species, PRDM6 is currently classified as a pseudo-methyltransferase that may interact with the methyltransferase G9a and corepressors to indirectly affect the methylation status of histone H3 lysine 9 and histone H4 lysine 20^6–8^.

Several PRDM proteins have been implicated in cellular processes related to stem cell maintenance or differentiation in the central nervous system^9^, but the role of PRDM6 in the developing and mature brain is not understood. Further, the role of PRDM6 expression in the etiology of Group 4 medulloblastoma or other brain cancers has not been tested. To assess if PRDM6 is a driver of Group 4 medulloblastoma, we set out to elucidate the functional impact and oncogenic potential of sustained PRDM6 expression in human neuroepithelial stem (NES) cells. Human NES cells are multipotent and retain the ability to give rise to cerebellar lineage cells, including granule neuron progenitors in cell culture and after orthotopic implantation in mice^10,11^. Because of their genetic stability and competence as cerebellar progenitors, NES cells are well-suited for studies of medulloblastoma development^11–14^.

## RESULTS

### PRDM6 expression and subcellular localization in human hindbrain neuroepithelial stem (NES) cells

We used human hindbrain neuroepithelial stem (NES) cells (**Figure 1A**), a representative of cerebellar progenitors that normally do not express PRDM6 (**Figure 1B**), to elucidate the molecular function and tumorigenic potential of PRDM6. We transduced NES cells with lentiviral vectors expressing V5 epitope-tagged human PRDM6 (hereafter referred to as “PRDM6 NES” cells) or the parental, empty lentiviral control vector (hereafter referred to as “EV NES” cells) (**Figure 1A**). qRT-PCR analysis and immunoblotting showed robust PRDM6 expression at the transcript and protein levels, respectively, in PRDM6 NES but not in EV NES cells (**Figure 1B** and **C**).

**Figure 1.**
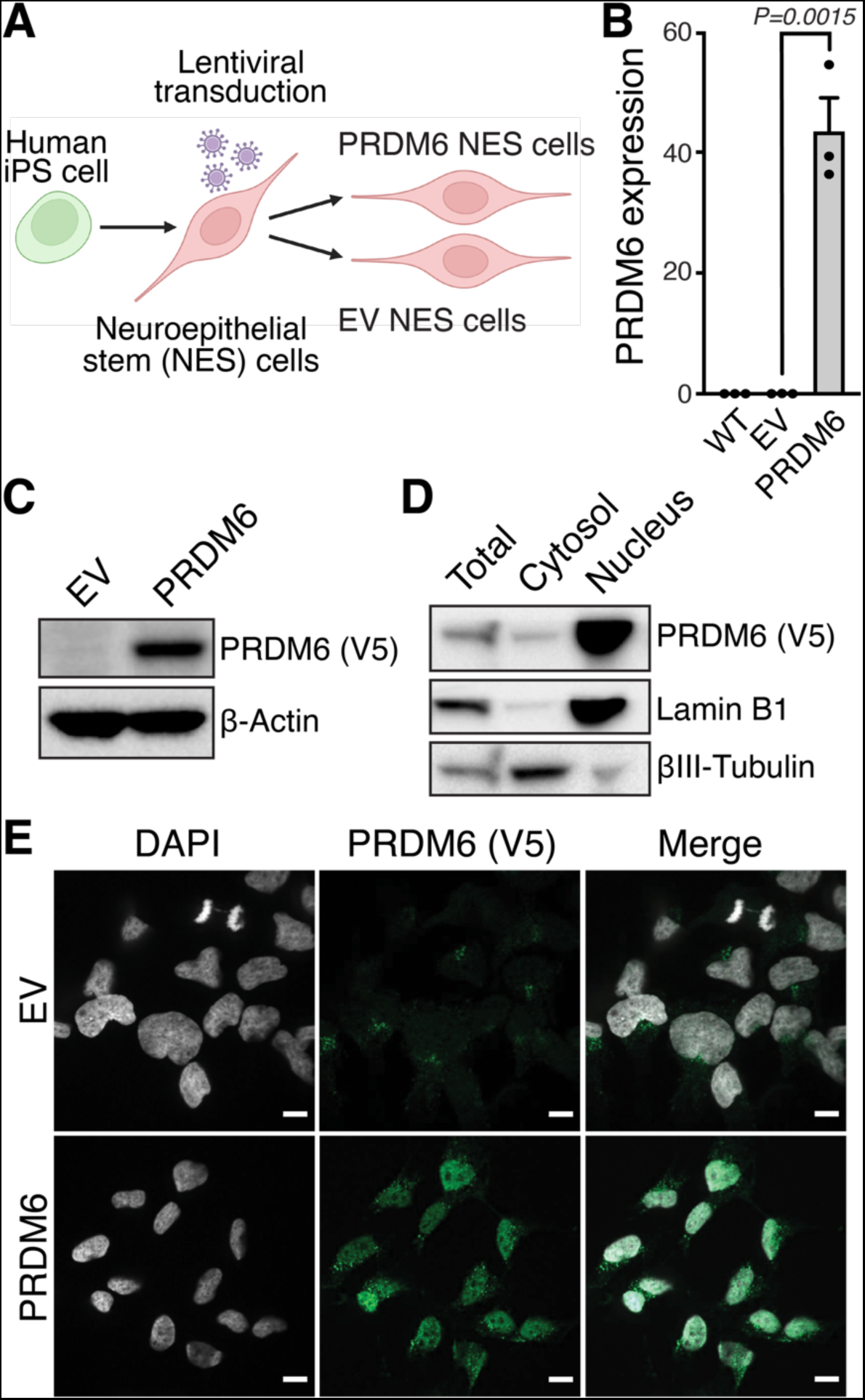
Expression and subcellular localization of PRDM6 in human neuroepithelial stem cells. **(A)** Schematic showing the generation of PRDM6-expressing NES cells from human iPS cells. **(B)** Wild-type (WT) NES cells do not express PRDM6. qRT-PCR analysis of *PRDM6* expression in wild-type, empty vector (EV), and PRDM6 expressing (PRDM6) NES cells. Expression levels are fold changes relative to *GAPDH*. Error bars denote SEM. *P*-values were determined by an unpaired, two-tailed *t*-test. **(C)** PRDM6 protein levels in EV and PRDM6 NES cells. Equal amounts of whole cell extract from empty vector (EV) or PRDM6-V5-transduced (PRDM6) NES cells were probed with anti-V5 antibodies to assess PRDM6 expression and antibodies to β-Actin as a loading control. **(D)** Subcellular fractionation of PRMD6 NES cells. Equal amounts of whole cell extract (total), cytosol, and nuclear extract from NES cells expressing V5-epitope-tagged, human PRDM6 were probed with antibodies against V5 and the indicated proteins of known subcellular localization. **(E)** Representative confocal microscopy images of anti-V5-immunostained EV and PRDM6 NES cells. Nuclei were counterstained with DAPI. Scale bars, 10 μm.

The subcellular localization of human PRDM6 has not been reported. To gain insight into its molecular function, we first assessed its subcellular localization in NES cells. Consistent with prior reports on mouse PRDM6^7,8^, subcellular fractionation (**Figure 1D**) and immunofluorescence microscopy (**Figure 1E**) experiments revealed a predominantly nuclear localization, indicating that PRDM6 functions in the nucleus in human NES cells.

### PRDM6 represses chromatin accessibility

Given the potential roles of other PRDM proteins in chromatin modification and gene expression^6^, we set out to elucidate the impact of human PRDM6 expression on chromatin accessibility. To this end, we performed a genome-wide analysis of chromatin accessibility by *Assay for Transposase-Accessible Chromatin using sequencing* (ATAC-seq) of PRDM6 and EV NES cells, respectively (**Figure 2**). First, we performed a qualitative occupancy analysis in PRDM6 and EV NES cells (*n*=3 biological replicates per group) to elucidate chromatin accessibility regions common or unique to each condition. This analysis revealed 11,076 open chromatin regions in PRDM6 NES cells, of which 10,844 (97.9%) were shared with EV NES cells (**Figure 2A**). In contrast, EV NES cells showed 35,273 open chromatin regions, of which 24,429 (69.26%) were absent in PRDM6 NES cells.

**Figure 2.**
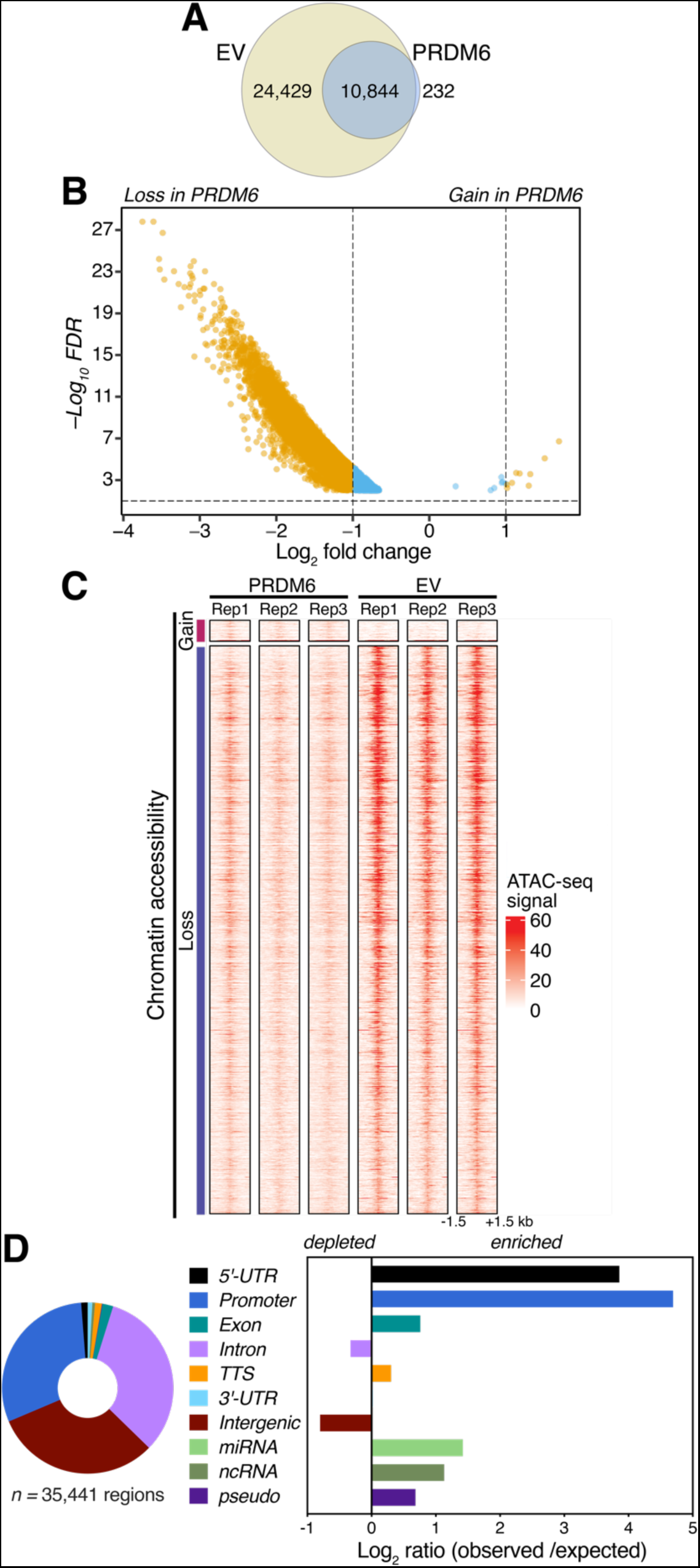
PRDM6 represses chromatin accessibility in neuroepithelial stem cells. **(A)** Venn diagram showing the numbers and overlap of regions with accessible chromatin in EV (yellow) and PRDM6 (blue) NES cells. **(B)** Volcano plot of results from a quantitative analysis of differential chromatin accessibility in PRDM6 *vs.* EV NES cells (*FDR* < 0.01). Regions with a *Log2* fold change ≥1 in chromatin accessibility are highlighted in yellow. Blue dots correspond to regions with significant (*FDR* <0.01) but <1 *Log2* fold change. **(C)** Heatmaps showing ATAC-seq signal within 1.5-kb of all sites with differential chromatin accessibility (*n*=35,441 sites, *FDR* < 0.01) across three replicate experiments (Rep1-3) in PRDM6 and EV NES cells. **(D)** *left,* Distribution of regions with differential accessibility (*FDR* < 0.01) across the indicated genomic annotations. *Right*, distribution of differential accessibility regions in PRDM6 NES relative to the expected genomic distribution.

Next, we performed a quantitative analysis of differential chromatin accessibility in PRDM6 *vs.* EV NES cells. In total, 35,441 regions showed differential chromatin accessibility between the conditions (false-discovery rate (*FDR*) ≤ 0.01). Strikingly, only 16 genomic regions (0.045%) were found to be more accessible, whereas 35,425 regions (99.95%) showed reduced chromatin accessibility upon PRDM6 expression, respectively. When considering only regions with ≥2-fold changes in chromatin accessibility, 24,298 regions showed reduced accessibility and only 9 regions showed increased accessibility in PRDM6 NES cells (**Figure 2B**). The very few genomic regions that gained accessibility upon PRDM6 expression showed only small increases in accessibility (**Figure 2B** and **C**).

Analysis of the genomic distribution of the 35,441 regions with differential chromatin accessibility between PRDM6 and EV NES cells revealed regions with altered accessibility not only in 5’-UTRs and promoters, but also in intronic and intergenic regions, and in other genome ontologies (**Figure 2D**, *left*). Differential accessibility regions were most enriched in 5’-UTRs, promoters, exons, miRNAs, ncRNAs, and pseudogenes (**Figure 2D**, *right*), indicating a predominant role of PRDM6 in those regions. However, the presence of differential accessibility regions in intronic and intergenic regions suggests that PRDM6 may also repress chromatin accessibility of enhancer regions located within intronic and intergenic regions.

Notably, gene ontology analysis of all regions with differential chromatin accessibility caused by PRDM6 overexpression in NES cells revealed a strong association with medulloblastoma (**Table S1**), suggesting that PRDM6 expression may prime NES cells for medulloblastoma development by affecting chromatin accessibility.

### PRDM6 expression deregulates gene expression in NES cells

As indicated above, regions with differential chromatin accessibility induced by PRDM6 expression were most strongly enriched in promoters and 5’-UTRs (**Figure 2D**, *right*). This prompted us to investigate the impact of PRDM6 on gene expression. Comparative RNA-seq analysis of PRDM6 and EV NES cells (*n*=3 biological replicates per group) revealed that the expression of a total of 1,588 genes was altered ≥2-fold (adjusted *P*-value < 0.01) (**Figure 3A**). Of these, 1,039 (65.43%) genes were upregulated and 549 genes (34.57%) were downregulated in PRDM6 NES cells. Gene ontology analysis of genes with ≥2-fold upregulation indicated a significant association with processes related to extracellular matrix (ECM) organization, ECM glycoproteins, collagens and proteoglycans (**Figure 3B**, *top*), whereas genes that showed ≥2-fold downregulation were most significantly associated with processes related to neuronal and synaptic processes (**Figure 3B**, *bottom*).

**Figure 3.**
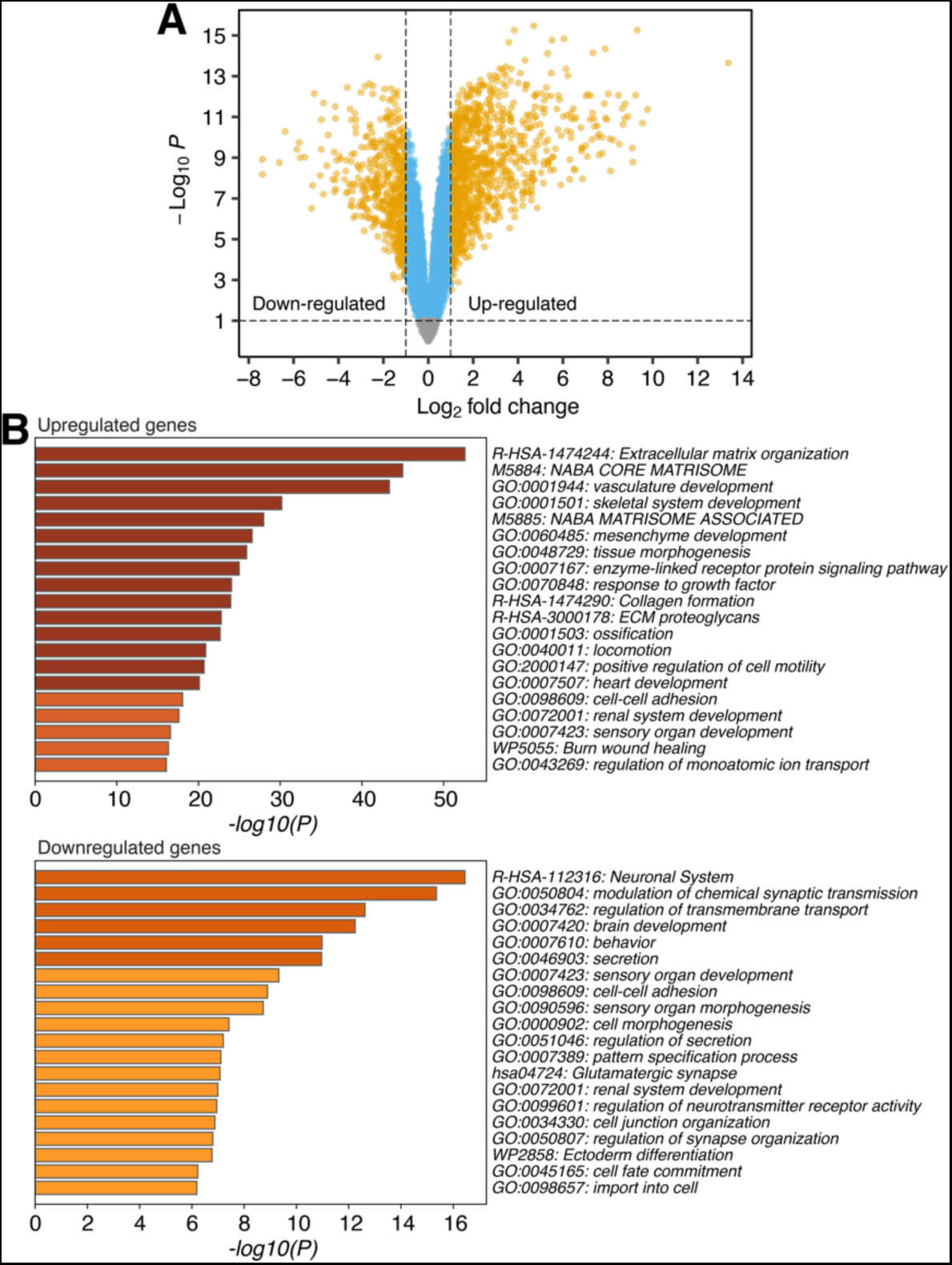
PRDM6 expression causes widespread gene expression changes in neuroepithelial stem cells. **(A)** Volcano plot of gene expression changes in PRDM6 *vs.* EV NES cells (*n*=3 biological replicates). Dots correspond to individual genes. Genes with significant (adjusted *P*-value <0.01) up- or downregulation (*Log2* fold change ≥1) are highlighted in yellow; genes with significant *Log2* fold changes of <1 are in blue. Gray dots indicate genes without significant changes (adjusted *P*-value >0.01). **(B)** Gene ontology (GO) analysis of upregulated (*top*) and downregulated (*bottom*) genes in PRDM6 NES cells. The bars correspond to significantly enriched GO terms and are colored by *P*-values in *log* base 10.

### PRDM6 affects transcription factor binding in open chromatin regions in NES cells

Chromatin accessibility modulates transcription by impacting the physical access of transcription factors and other regulatory factors, thus affecting cellular function and identity. To assess the potential mechanistic implications of the observed changes in gene expression and chromatin accessibility between PRDM6 and EV NES cells, we analyzed all genomic regions with differential accessibility for known transcription factor binding motifs *via HOMER*. This analysis revealed significant enrichment of binding motifs for transcription factors known to be involved in the regulation of genome organization, development, cell fate, and tumor suppression (**Table S2**).

To further interrogate the implications of the PRDM6-driven, altered chromatin accessibility landscape in NES cells, we performed footprinting of transcription factor binding in all open chromatin regions *via Transcription factor Occupancy prediction by Investigation of ATACseq signal* (*TOBIAS*)^15^. Whereas *HOMER*-based motif enrichment analysis can reveal enriched transcription factor binding motifs within genomic regions, *TOBIAS* quantifies the occupancy of transcription factor motifs and infers which transcription factor is bound according to the sequence of the actual footprint. Footprinting analysis of all open chromatin regions revealed 841 transcription factor binding sites (**Figure 4** and **Table S3**), of which 73 showed a differential binding score of ≥0.1 (*P<0.0001*) in PRDM6 *vs.* EV NES cells. Among these, 15 transcription factors (20.55%) showed more binding in PRDM6 NES cells, whereas 58 transcription factors (79.45%) showed less binding in PRDM6 NES cells (**Figure 4A** and **Table S3**). The binding of RFX and SOX family was most strongly impaired by PRDM6 expression in NES cells (Figure 4B). Notably, RFX transcription factors, including RFX1^16^, have been implicated in tumor suppression^17^, suggesting that reduced binding of these transcription factors in the presence of PRDM6 may have oncogenic potential in NES cells. Moreover, the binding of other transcription factors with roles in genome organization, development, and cell fate was affected (**Table S3**). Together, these findings support the notion that PRDM6 affects the developmental and oncogenic potential of NES cells by altering transcription factor binding.

**Figure 4.**
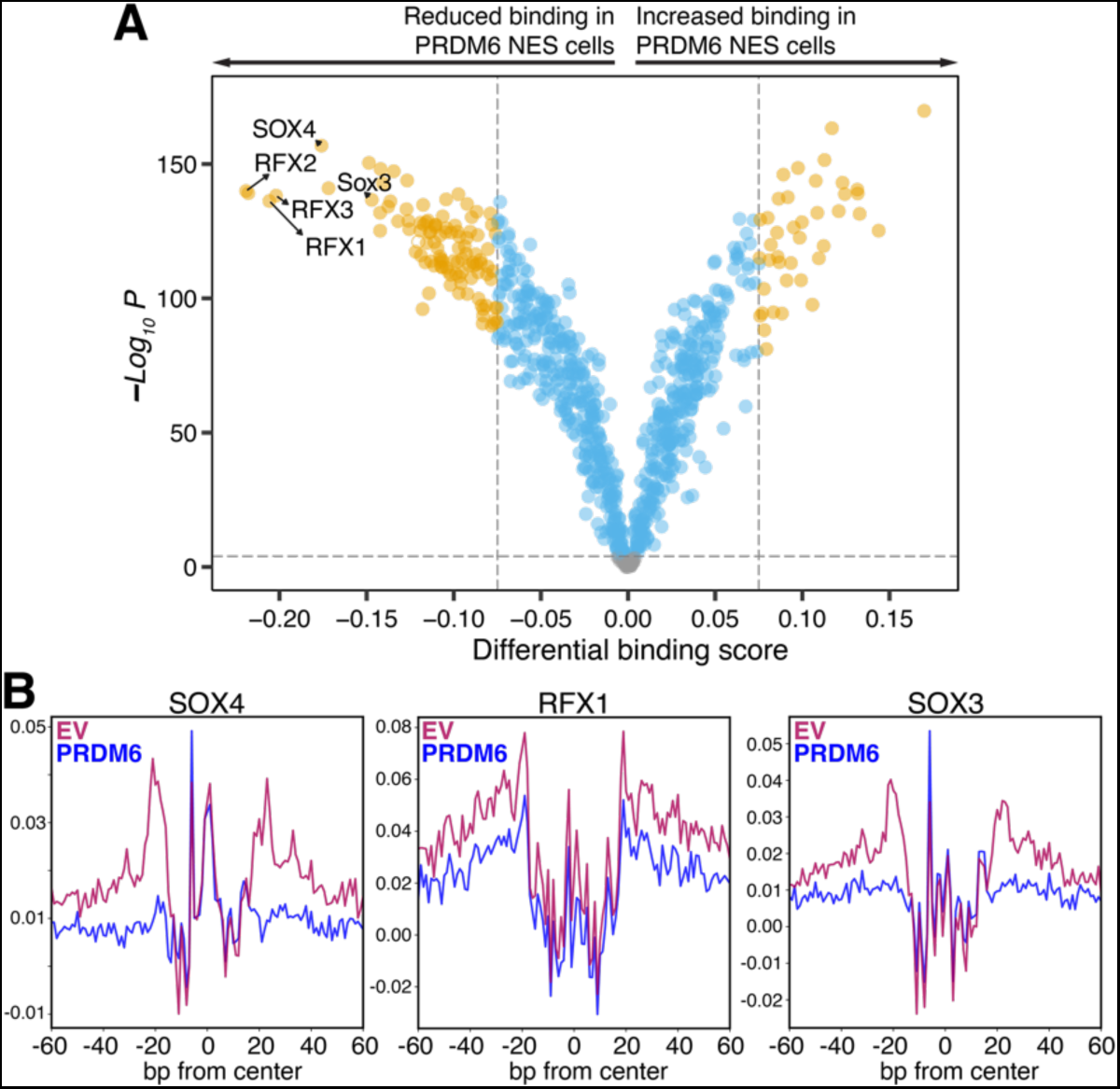
Impact of PRDM6 on transcription factor binding in open chromatin regions. **(A)** Volcano plot showing the quantification of differential transcription factor binding based on digital footprinting analysis of *JASPAR* transcription factor motifs (*n*=841) in open chromatin regions of PRDM6 and EV NES cells. Transcription factor motifs with significantly differential binding scores (*P*<0.0001) are indicated in blue; transcription factor motifs with high (≥0.075) changes in differential binding score (*P*<0.0001) are highlighted in orange. Gray dots represent transcription factor motifs without significantly altered binding scores (*P*≥0.0001). Footprinting was performed in *n*=3 biological replicates per group. **(B)** Aggregate plots of footprinting signal across all SOX4, RFX1, and SOX3 motifs in EV (purple) and PRDM6 (blue) NES cells.

### Genome-wide mapping of PRDM6 binding reveals colocalization with genes in H3K27me3-rich regions

To further elucidate the molecular function of PRDM6, we used *Cleavage Under Targets and Release Using Nuclease* (CUT&RUN)^18^ to map the genome-wide distribution of PRDM6 in NES cells. In the absence of reliable antibodies against endogenous human PRDM6, we used V5 antibodies for CUT&RUN in EV and PRDM6 NES cells. We used the *nf-core* best-practices CUT&RUN analysis pipeline^19,20^ to identify peak regions and performed differential peak calling by using EV NES cells as controls. This resulted in the detection of 1,083 peaks in PRDM6 NES cells, which were absent in EV NES cells (**Figure 5B**). PRDM6 binding sites in the NES genome were enriched in 5’-UTRs, promoters, and other genomic features but were depleted in intergenic regions (**Figure 5A**). 952 of 1,083 PRDM6 peaks (87.90%) overlapped or were located within two kb of an annotated gene, which implicates PRDM6 in gene regulation (**Table S4**).

**Figure 5.**
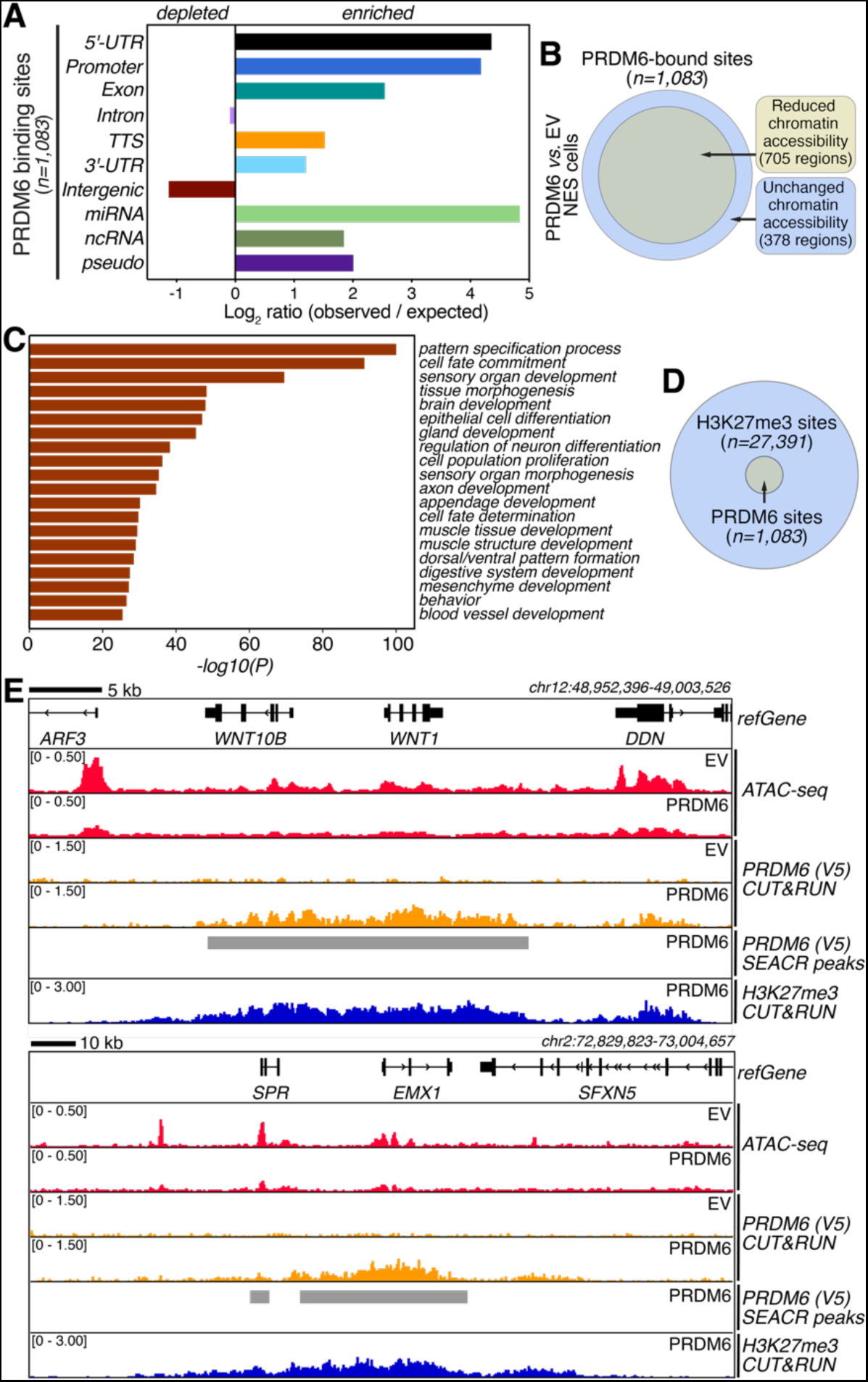
Elucidation of genome-wide PRDM6 binding in neuroepithelial stem cells. **(A)** Genomic distribution of PRDM6 binding sites determined by CUT&RUN **(B)** Venn diagram showing the overlap of PRDM6-bound sites with regions showing altered chromatin accessibility (*FDR* < 0.01). **(C)** Gene ontology (GO) analysis of genes overlapping with or located within two kb of a PRDM6 binding site. Bars are colored by *P*-values in *log* base 10 and indicate significantly enriched GO terms. **(D)** Venn diagram showing the overlap of PRDM6-bound sites with H3K27me3-marked regions identified by CUT&RUN in PRDM6 NES cells. **(E)** Examples showing PRDM6 enrichment in relation to H3K27me3-marked regions. Bins per Million (BPM)-normalized CUT&RUN and ATAC-seq signals are shown. SEACR peak calling results for PRDM6 CUT&RUN signal are indicated by gray bars. Genomic region coordinates and *refGene* annotations are shown.

705 of 1,083 (65.10%) PRDM6-bound genomic regions overlapped with regions showing significantly reduced chromatin accessibility in PRDM6 NES cells (**Figure 5B**), suggesting a predominantly repressive effect of PRDM6 binding to chromatin. However, given the relatively small number of PRDM6 binding sites detected by CUT&RUN, most regions with differential accessibility did not show PRDM6 binding. There are at least two, not mutually exclusive, interpretations of this finding: first, most of the effects of PRDM6 on chromatin accessibility occur through indirect mechanisms; second, binding of PRDM6 to chromatin is indirect or very transient and hence difficult to detect by CUT&RUN, which results in underestimation of the true number of PRDM6 binding sites in the genome.

Gene ontology analysis of genes overlapping with or located within two kilobases (kb) of a PRDM6 binding site revealed a strong enrichment of terms related to developmental and neural processes (**Figure 5C**). Further, *Molecular Signatures Database* (MSigDB)^21^ analysis, suggested a significant association with processes related to histone H3 lysine 27 trimethylation (H3K27me3) (**Table S5**). To follow up on this latter observation, we determined the H3K27me3 landscape in PRDM6 NES cells *via* CUT&RUN. Bioinformatic analysis revealed 27,391 H3K27me3 binding sites. Notably, all 1,083 PRDM6 binding sites overlapped H3K27me3 binding sites (**Figure 5D**), suggesting a functional interplay between H3K27me3-related processes and PRDM6. In the latter regard, the overlap of PRDM6 binding sites with H3K27me3 regions (**Figure 5D and E**) is consistent with the predominantly repressive effect of PRDM6 binding reported above. We did not, however, detect changes in global H3K27me3 levels in PRDM6 *vs*. EV NES cells (**Figure S1**). This is perhaps not surprising given that we detected PRDM6 binding in only approximately 3.95% of H3K27me3 regions (**Figure 5D**). Nevertheless, the finding that PRDM6 binding completely overlaps with chromatin marked by H3K27me3, which affects spatiotemporal control of gene expression in stem cells, further implicates PRDM6 as a modulator of chromatin structure and gene expression in NES cells.

### PRDM6 expression in NES cells promotes medulloblastoma formation

Having revealed the molecular impact of PRDM6 expression in NES cells, we asked whether the observed changes in chromatin landscape and gene expression establish a cellular environment prone to tumorigenesis. To test the tumorigenic potential of PRDM6 in NES cells, we performed *in vitro* proliferation assays and orthotopic intracranial implantation experiments in mice (**Figure 6A**). *In vitro,* PRDM6 NES cells showed no change in proliferation rate when compared to NES cells without PRDM6 expression (**Figure S2A**). Next, we orthotopically implanted 3 × 10^5^ cells into the cerebellum of adult, female NSG mice and monitored tumor growth for a year. About a third (8/30) of mice implanted with PRDM6-expressing NES cells developed tumors with a latency of 158 ± 16 days (mean ± SEM) after transplantation (**Figure 6B**). In contrast, implantation of NES cells without PRDM6 expression (EV) did not cause tumor formation (0/15) within a year of implantation, the endpoint in our study. These findings indicate that PRDM6 expression can indeed establish a pro-tumorigenic milieu.

**Figure 6.**
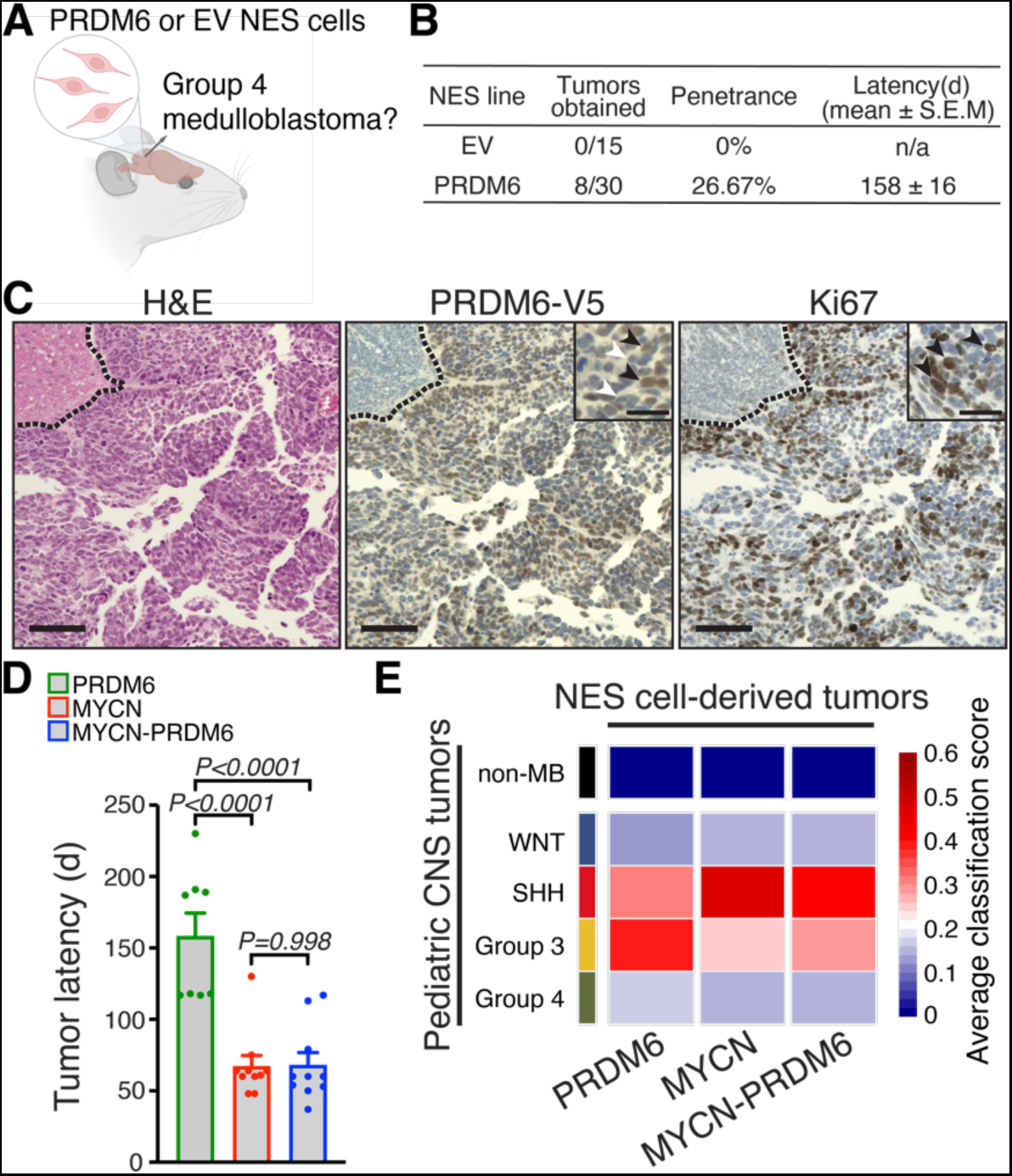
PRDM6 expression in neuroepithelial stem cells induces medulloblastomas in mice. **(A)** Schematic of orthotopic human NES transplantation into the mouse cerebellum. **(B)** Outcomes of PRDM6 or EV NES cell transplantations in mice. d, days post-injection. **(C)** H&E staining of representative PRDM6 NES-derived tumor (*left*). Immunohistochemistry staining of PRDM6 *via* the V5 epitope tag (*middle*) and Ki67 (*right*) in PRDM6 NES-derived tumor. Dashed lines denote tumor margins. Main scale bars correspond to 100 μm; scale bars in insets correspond to 25 μm; black arrowheads in the middle panel inset denote anti-V5-reactive (PRDM6-expressing) nuclei; white arrowheads indicate PRDM6-negative nuclei; black arrowheads in the right panel inset denote Ki67-positive, proliferating cells. **(D)** Latency of tumor development after implantation of PRDM6-, MYCN- or MYCN-PRDM6 NES cells into the cerebellum of mice. Error bars represent SEM. *P-*values were determined by one-way ANOVA with Tukey’s *post hoc* correction. **(E)** Clustering of tumors derived from PRDM6-, MYCN-, or MYCN-PRDM6 NES cells with human medulloblastomas and pediatric, non-medulloblastoma (non-MB) CNS tumors.

Next, we performed a histopathological analysis of tumors derived from PRDM6 NES cells. Microscopic examination of hematoxylin and eosin (H&E)-stained tumor tissue revealed a highly cellular neoplasm with mitotic activity and apoptotic bodies (**Figure 6C***, left panel,* **Figure S3**). Tumors lacked glial components and showed an overall morphology reminiscent of primitive neuroectodermal tumors, which is consistent with human medulloblastoma histopathology (**Figure 6C**, *left panel*). Increased mitotic activity was further supported by strongly-positive Ki67 immunostaining (**Figure 6C**, *right panel*). Moreover, immunostaining for PRDM6 *via* detection of its V5 epitope tag showed that NES-derived tumors had maintained PRDM6 expression, suggesting that PRDM6 may function in both tumor initiation and maintenance (**Figure 6C**, *middle panel*).

Because a subset of Group 4 medulloblastomas shows high expression of PRDM6 in conjunction with amplification of the *MYCN* oncogene^1,4^, we decided to investigate potential synergies between PRDM6 and MYCN expression on tumor development from NES cells. Notably, MYCN expression alone is sufficient to drive tumor formation from NES cells, resulting in SHH medulloblastoma^11^. We generated human NES cells with either MYCN overexpression (“MYCN” cells), or combined MYCN and PRDM6 overexpression (“MYCN-PRDM6” cells), respectively (**Figure S2B** and **C**). MYCN-overexpressing NES cells showed significantly faster proliferation *in vitro*, irrespective of PRDM6 expression (**Figure S2D**).

To evaluate the impact of combined PRDM6 and MYCN overexpression on tumor formation, we performed orthotopic transplantation experiments in mice. Mice implanted with MYCN- or MYCN-PRDM6 NES cells showed 100% penetrance of tumor growth in both groups (**Figure S2E**). Latencies of tumor development from MYCN- *vs.* MYCN-PRDM6 NES cells were not different (67 ± 7 days and 68 ± 8 days (median ± SEM), respectively; **Figure S2E**). Notably, these latencies were significantly shorter than those observed after the transplantation of PRDM6 NES cells (**Figure 6D**). Overall, these findings suggest that PRDM6 does not modulate the tumorigenic effects of MYCN overexpression in NES cells.

To molecularly characterize tumors arising from PRDM6- or MYCN-PRDM6 NES cells, we performed RNA-seq analysis. We then carried out a comparative gene expression clustering analysis with medulloblastomas and other pediatric central nervous system (CNS) tumors (*i.e.*, “non-medulloblastoma” tumors) from human patients (**Figure 6E**). For medulloblastoma subgroup clustering, we used a classifier trained on 1,112 medulloblastomas (WNT, SHH, Group 3, Group 4) and 1,473 primary, non-medulloblastoma tumors corresponding to 10 distinct CNS tumor types. Tumors derived from PRDM6 NES cells clustered most closely with Group 3 medulloblastomas and showed no similarity with pediatric non-medulloblastoma CNS tumors (**Figure 6E**). In contrast, tumors derived from MYCN NES cells clustered most closely with SHH medulloblastomas, consistent with previous findings (F**igure 6E**)^11^. Tumors derived from MYCN-PRDM6 NES cells clustered with SHH medulloblastomas as well, suggesting that MYCN has a dominant effect on tumor phenotype (**Figure 6E**). These latter findings suggest that PRDM6 expression does not modulate the onset, progression, or molecular phenotypes of tumors derived from MYCN-overexpressing NES cells.

Together, our findings show that PRDM6 expression in NES cells has oncogenic potential and promotes medulloblastoma formation. However, we conclude that additional factors are required for Group 4 medulloblastoma formation as PRDM6 NES cell-derived tumors most closely resemble Group 3 medulloblastoma.

## DISCUSSION

### Molecular function of PRDM6

The PRDM family is implicated in mammalian neurodevelopment and control of cell proliferation in cancer^22^ but the role of PRDM6 in these processes is unclear. We found that human PRDM6 functioned in the nucleus, exerting a strongly repressive effect on chromatin accessibility in hindbrain neuroepithelial stem (NES) cells. Moreover, we found a complex dysregulation of gene expression patterns in PRDM6 NES cells, with substantial downregulation and upregulation of genes. These findings are consistent with the notion that human PRDM6 exists in protein complexes containing both transcriptional activators and repressors. Such binary roles have been shown for PRDM1, another PRDM family member, which recruits the histone demethylase KDM4a to remove repressive H3K9me3 marks^23^ and also interacts with histone deacetylases to repress transcription^24^. Moreover, mouse PRISM/PRDM6 has been implicated in both transcriptional activation and repression^8^, consistent with our findings in human NES cells. It will be important to elucidate the composition of human PRDM6-containing protein complexes to fully reveal the mechanisms by which PRDM6 affects chromatin structure and gene expression. In this regard, we found that PRDM6 bound to H3K27me3-rich regions. H3K27me3 affects the spatiotemporal control of gene expression in complex ways, so it will be important to interrogate interplays between H3K27me and PRDM6 in NES cells and in medulloblastoma.

### Oncogenic potential of PRDM6

Sequencing of medulloblastomas has revealed many molecular aberrations. A key challenge is to distinguish “driver” from “passenger” alterations. Our findings that PRDM6 expression in hindbrain neuroepithelial stem cells promotes medulloblastoma indicate that PRDM6 can indeed function as a medulloblastoma driver.

Several, not mutually exclusive reasons may underlie why PRDM6 NES cell-derived medulloblastomas do not recapitulate Group 4 medulloblastomas. Additional factors such as a specific cell-of-origin, co-occurring genomic alterations, or the specific microenvironment of the developing cerebellum may contribute. Correspondingly, it will be important to test the oncogenic impact of PRDM6 expression in other cerebellar progenitors, *e.g.,* in unipolar brush cells^1,25^, ideally in the immature embryonic hindbrain^1,2^, or in the context of other genomic alterations that occur in PRDM6-expressing Group 4 medulloblastomas^4^.

Our observations may provide clues as to how PRDM6 expression promotes medulloblastoma formation from NES cells. Our footprinting analysis shows significantly altered binding of transcription factors to open chromatin regions in PRDM6 NES cells. These include RFX1, a transcriptional repressor of tumor suppressors, and several other RFX transcription factors^16^. Notably, multiple RFX1 target genes, including the oncogene *FGF1*, *COL1A1*, and *COL1A2*, are upregulated in PRDM6 NES cells. Moreover, RFX1 dimerizes with RFX2 and RFX3^26^, which also show reduced binding in PRDM6 NES cells (**Figure 4**). These findings may provide a starting point for how PRDM6 establishes a pro-oncogenic environment. In the latter context, we find that PRDM6 expression results in substantial upregulation of processes related to extracellular matrix (ECM) biology (**Figure 3B**). The ECM is important for cell adhesion, growth factor binding, and intracellular signaling. Importantly, tumors use ECM remodeling to promote tumorigenesis and metastasis^27^. In future studies, it will be important to address the functional implications of PRDM6-driven ECM alterations as it may yield insights into the tumorigenic effects of PRDM6.

Given the oncogenic potential of PRDM6 we report here, its inhibition may be of benefit in PRDM6-expressing Group 4 medulloblastomas. PRDM6 expression is shut off postnatally^22,28^, which would make it an attractive drug target as PRDM6-selective inhibition should not affect the homeostasis of normal tissues.

## METHODS

### Cell culture

Human neuroepithelial stem (NES) cells were generated from human iPS cells (WTC10^29^) as previously described^11,12^. Experimental work involving iPS cells was approved by the UCSF Stem Cell Research Oversight Committee. All cell lines used here were regularly screened for mycoplasm and were mycoplasm-negative by PCR-based testing. NES cells were cultured on poly-L-ornithine/laminin-coated wells in NES cell medium (DMEM/F-12 with GlutaMAX, 1× N2 supplement, 0.05× B27 without vitamin A supplement, 1.6 g/L Glucose, 20 ng/mL EGF, 20 ng/mL FGF2) in a humidified 37 °C incubator with 5% CO_2_. NES cells were fed daily, passaged every three to four days using TrypLE Express, and plated at 5-6 × 10^5^ cells per well of a 6-well plate in 2 mL of NES cell medium. For lentiviral transductions, NES cells were plated on poly-L-ornithine/laminin-coated wells, and PEG-concentrated lentiviruses were added to the cells. Lentiviruses were washed out 24 h after transduction, cells were expanded, and subjected to either antibiotic selection starting 72 h after transduction or sorted by flow cytometry.

### Mice

All studies followed ethical regulations established by the Institutional Animal Care and Use Committee and Institutional Biosafety Committee at the University of California, San Francisco. The reporting in this manuscript follows the ARRIVE guidelines^30^. 8-12 week-old female NSG (*RRID:IMSR_JAX:005557*) mice were used for transplantation experiments. For transplantation, 3 × 10^5^ cells in 3 μL of NES cell medium were implanted per mouse. Implantations were performed using a stereotactic frame, starting from lambda 2 mm right, 2 mm down, and 2 mm deep. Mice were euthanized at endpoint, which was either signs of tumor growth (*e.g.*, hunched back, weight loss, head tilt, *etc*.) or one year after transplantation. For monitoring tumor growth *in vivo*, mice were subjected to weekly or bi-weekly luciferase-based bioluminescence imaging.

### Lentiviral constructs

The coding sequence of human PRDM6 (*NCBI* Reference Sequence NM_001136239.4) was fused to a C-terminal V5-tag separated by a 2× glycine linker and inserted into a lentiviral vector to generate pCDH-Lenti-CAGGS-hPRDM6-2×Gly-V5-EF1-Luc2-Blast. The lentiviral MYCN expression plasmid pCDH-CAG3×FLAG-MYCN-mScarlet-Luciferase was previously published^11^.

### Protein extraction and immunoblotting

Protein extracts were prepared using RIPA buffer (150 mM NaCl, 1% IGEPAL CA-630, 0.5% sodium deoxycholate, 0.1% SDS, 50 mM Tris, pH 8.0; Millipore Sigma) supplemented with protease inhibitors (*cOmplete* protease inhibitor cocktail; Roche) and quantified by BCA assay. Proteins were separated through a 4-12% (w/v) Bis-Tris SDS poly-acrylamide gel and transferred onto nitrocellulose membranes. Blots were incubated with primary antibodies in 5 % (w/v) BSA in TBS-T at 4 °C overnight, washed in TBS-T, and incubated with secondary antibodies in 5 % (w/v) BSA in TBS-T for 2 h at room temperature. Membranes were washed in TBS-T and developed using ECL chemiluminescence reagent.

### Subcellular fractionation

Whole-cell, nuclear, and cytosolic extracts from NES cells were prepared by using the Nuclear Extract Kit (Active Motif) as per the manufacturer’s instructions. In brief, cells were scraped into ice-cold PBS supplemented with protease inhibitors (*cOmplete* protease inhibitor cocktail, Roche) and phosphatase inhibitors, sedimented by centrifugation, and resuspended in hypotonic lysis buffer (Nuclear Extract Kit, Active Motif). Cytosolic and nuclear fraction were separated by centrifugation (14,000 × g for 10 min at 4 °C). Nuclei were extracted in Complete Lysis Buffer (Active Motif AM1 Lysis buffer containing 1 mM DTT and protease inhibitors). Whole-cell fractions were prepared by direct extraction of cells in Complete Lysis Buffer. Subcellular fractions were normalized by volume and analyzed by immunoblotting.

### Histone extraction

Histones were extracted by H_2_SO_4_ extraction as previously described^31^. Briefly, cell pellets were lysed in hypotonic lysis buffer (10 mM Tris-HCl, pH 8.0, 1 mM KCl, 1.5 mM MgCl_2_, 0.5 % (v/v) NP40, 1 mM DTT, *cOmplete* protease inhibitor cocktail). Nuclei were sedimented at 600 × g for 5 min at 4 °C. The nuclear pellet was incubated in 0.4 N H_2_SO_4_ for 16-18 h at 4 °C. Histones were precipitated with trichloroacetic acid, washed with acidified acetone, resuspended in water, and processed for immunoblotting.

### Cell proliferation assays

*In vitro* proliferation assays of NES cells were performed using either *Cell Titer Glo* (Promega) or *CyQUANT NF* (Thermo Fisher) according to the manufacturer’s instructions. In brief, 5 × 10^3^ NES cells were plated in 100 μL of NES media in triplicates for each condition. Proliferation was measured at timepoint day 0 (d0, *i.e.*, 2 h after cell plating) and every 24 h thereafter, for seven days. Levels of proliferation are represented as fold changes over d0 values.

### Histology, immunohistochemistry, and immunofluorescence microscopy

Immunohistochemistry was performed on formalin-fixed, paraffin-embedded tissue. In brief, paraffin blocks were cut into 5-μm thick sections, deparaffinized with xylene substitute, and rehydrated with decreasing concentrations of ethanol in water. For antigen retrieval, tissue slides were incubated in sodium citrate buffer (pH 6.0), followed by quenching in 0.3% (v/v) hydrogen peroxide. Tissue was permeabilized in blocking buffer (0.4% (v/v) Triton-X-100, 10% (v/v) normal goat serum in phosphate-buffered saline (PBS)). Primary antibodies were applied for 16-18 h at 4 °C in a humidified chamber. Sections were incubated with biotinylated secondary antibody for 1 h at room temperature in a humidified chamber. For signal amplification, slides were incubated with Vectastain ABC reagent and color development was achieved by diaminobenzidine tetrahydrochloride solution. Sections were counterstained with hematoxylin, dehydrated through ethanol and xylene substitute, and cover-slipped using mounting medium. Stained sections were imaged using a digital microscope (Keyence BZ-X) and BZ-X Keyence software at 40× and 100× magnifications. For immunofluorescence staining, NES cells were cultured on poly-L-ornithine/laminin-coated coverslips. NES cells were fixed in 4% (w/v) paraformaldehyde in PBS for 15 min at room temperature. Quenching was performed in 50 mM NH_4_Cl in PBS, followed by permeabilization in 0.5% (v/v) Triton-X-100 in PBS. Coverslips were incubated in blocking buffer (5% (v/v) goat serum, 0.1% (v/v) Tween in PBS) for 1 h at room temperature, followed by a 16-18-h incubation at 4 °C with V5 primary antibody (Invitrogen) diluted 1:50 in blocking buffer. Coverslips were washed in PBS and incubated in blocking buffer containing fluorescent secondary antibodies for 1 h at room temperature. Coverslips were washed in PBS and mounted with Fluoromount G/DAPI. Imaging was performed using a Zeiss Spinning Disk confocal microscope.

### ATAC-seq

Preparation of ATAC-seq libraries and sequencing was performed as described^32^, with slight modifications. In brief 1 × 10^5^ cells per sample were harvested and lysed in ATAC-seq lysis buffer (10 mM Tris-HCl (pH 7.4), 10 mM NaCl, 3 mM MgCl_2_, 0.1 % (w/v) IGEPAL CA-630). The transposition reaction was performed using Tagment DNA TDE1 Enzyme and Buffer Kit (Illumina) and 4.7 µL TDE1 Tagment DNA enzyme per sample in a 50-µL total reaction volume. Zymo DNA Clean and Concentrator Kit was used for cleanup of the transposition reaction. Quantitive PCR and limited-cycle PCR was performed using NEBNext Ultra II master mix (New England Biolabs), SYBR™ green and 0.5 µM of sample-specific P5/P7 barcoded primer mix (New England Biolabs). Bead clean-up and purification was performed using SPRI beads (Beckman Coulter). Agilent TapeStation 4200 was used to check for quality of sequencing libraries. To support standardization and reproducibility, we used the *nf-core* framework best-practice ATAC-seq analysis pipeline (*nf-core/atacseq* v1.2.1) for sequence processing and data analysis^19,33^. In brief, adapters were trimmed by TrimGalore (v0.6.4_dev)^34^ and adapter-trimmed reads were mapped to the *hg38* reference assembly by BWA (v0.7.17)^35^. Duplicates were marked and removed by Picard (v2.23.1)^36^. ATAC-seq peaks were called by MACS2 (v2.2.7.1)^37^ and differential analysis of ATAC-seq peaks present in all three biological replicates per group (PRDM6 or EV NES cells) was performed by DiffBind (v3.6.5)^38,39^ and DESeq2^40^. Transcription factor occupancy analysis was performed by TOBIAS (v0.15.1)^15^. Correction for Tn5 transposase insertion bias (*ATACorrect*) and footprint scoring (*ScoreBigwig*) were performed as described^15^. Footprint scores were matched to the *JASPAR 2022 CORE vertebrates (non-redundant)* database^41^ and differential binding scores were calculated by *BINDetect*^15^.

### CUT&RUN

Preparation of CUT&RUN libraries was done with the CUT&RUN assay kit (Cell Signaling Technology). Briefly, 2.5 × 10^5^ NES cells per sample were fixed with 0.1% formaldehyde in PBS for 2 min and quenched with 1 M glycine for 5 min at room temperature. Cells were washed in 1× wash buffer containing spermidine and protease inhibitor cocktail, and incubated in antibody binding buffer containing spermidine, proteinase inhibitor cocktail, digitonin, and 10 μl of activated concanavalin A beads per sample for 16-18 h at 4 °C. Per CUT&RUN reaction, 0.2 µg of H3K27me3 antibody (clone C36B11, #9733, Cell Signaling Technology) or 0.96 µg of V5 antibody (clone D3H8Q, #13202, Cell Signaling Technology) was used. pAG-MNase in 1× wash buffer containing spermidine, protease inhibitor cocktail, and digitonin was bound to the antibody for 1 h at 4 °C. pAG-MNase was activated by adding 3 μL calcium chloride per sample and incubated for 30 min at 4 °C. Digestion was stopped by adding 1× stop buffer (digitonin solution, RNase A, and 50 pg *S. cerevisiae* spike-in DNA (Cell Signaling Technology) for normalization) and samples were incubated for 10 min at 37 °C. Samples were subjected to SDS and proteinase K treatment for 2 h at 65 °C. Zymo DNA Clean and Concentrator (D4034) was used to clean up the samples. NEBNext Ultra II DNA Library Prep Kit for Illumina (E7645L) was used for library preparation. Library clean-up and size selection was done by using SPRI beads (103778-488, Bulldog Bio). Libraries were quantified and analyzed by Qubit and Tapestation 4200 (Agilent) analysis and sequenced for 38 cycles in paired-end mode on a Nextseq500 (Illumina). For data analysis, we used the nf-core framework CUT&RUN analysis pipeline (v3.0; https://github.com/nf-core/cutandrun/tree/3.0)^19,20^. In brief, adapter-trimmed reads (TrimGalore v0.6.6) were mapped to the human (*hg38*) and spike-in (*S. cerevisiae*, *sacCer3*) genome reference assemblies by Bowtie2 (v2.4.4). Duplicates were marked by Picard (v2.27.4). Reads were normalized against spike-in. CUT&RUN peaks were called by SEACR (v1.3) using stringent settings, and consensus peaks were identified as peaks that were present in both replicates per condition. The proximity of peaks to genes was assessed by T-Gene (v5.5.3)^42^.

### RNA isolation and quantitative RT-qPCR

RNA was extracted using an RNA extraction kit (Zymo Research). 500 ng of RNA was converted to cDNA using the High-Capacity cDNA Reverse Transcription Kit (Thermo Fisher) and the following settings: 25 °C for 10 min, 37 °C for 2 h, and 85 °C for 5 min. qPCR was performed using PowerUp SYBR green (Thermo Fisher) with the following settings: 95 °C for 2 min, 40 cycles [95 °C for 15 s and 60 °C for 1 min]. Gene expression was normalized to *GAPDH*.

### RNA-seq

Total RNA was extracted from cultured cells or flash frozen tumor tissue using RNA extraction kit (Zymo Research). RNA quality was checked using Agilent TapeStation 4200. Samples were processed for library preparation (polyA enrichment, Illumina TruSeq stranded mRNA kit) and sequenced (NovaSeq PE100 or PE150; 62-112 million total reads per sample). Raw RNA-seq data was pre-processed using *HTStream* (https://github.com/s4hts/HTStream). Pre-processed reads were aligned using STAR (v2.7.10, *GRCh38*). Differentially expressed genes (*Log2* fold change ≥1, adjusted *P*-value <0.01) were identified by *edgeR* and *limma-voom* analysis^43^.

### Classification of NES-derived tumors

Raw RNA-seq reads were processed to remove low-quality sequences and adapter sequences. To discriminate between human and mouse reads, *disambiguate* was used to filter out reads with a higher mapping quality score in the mouse genome compared to the human genome^44^. Human-specific reads were mapped to *GRCh38/hg38* using STAR to generate *Fragments Per Kilobase of transcript per Million* mapped reads (FPKM) values^45^. To quantify the resemblance of NES-derived tumors with medulloblastoma subgroups, we trained a machine-learning classification model on a large series of primary medulloblastoma tumors. Specifically, two published gene expression microarray series of medulloblastomas, with medulloblastoma subgroup determined by DNA methylation, were obtained (GEO GSE85218) and (EGAS00001001953). After removing duplicate samples this resulted in 1,112 unique expression profiles that underwent Robust Multi-array Average normalization. Because gene sets are more robust to platform differences than individual genes, single-sample gene set enrichment (ssGSEA) scores were derived by using all *MSigDB* gene sets that contained ≥80% mouse–human orthologs and ≥15 and ≤400 genes^46^. A random forest classification model was then trained using a 70-30 training-test split of the medulloblastoma expression profiles, using the gene set enrichment scores as features to predict medulloblastoma subgroup. Gene sets with a scaled feature importance score >8 were used to build the final classification model resulting in a total of 409 features (test set accuracy = 0.956). This model was used to predict the subgroup class probabilities for all NES-derived tumors. The same workflow was utilized to create a binary machine learning model to classify if tumors were medulloblastoma. The dataset used to train the non-medulloblastoma model consisted of 1,473 primary CNS tumors, after filtering undefined entities, encompassing 10 distinct CNS tumor types obtained from the R2 pediatric genome portal (accession “Tumor Brain (DKFZ-public)-Kool-1678-MAS5.0-u133p2”). Using a 70-30 training-test split, the non-medulloblastoma model achieved a test set accuracy of 0.987.

### Motif, genome ontology, and gene ontology analyses

Genome annotations were determined by *HOMER* version 4.11.1^47^ using *HOMER hg38* accession and ontology information with ‘*annotatePeaks.pl* –annStats’ and default settings. *HOMER*-based motif analysis used *findMotifsGenome.pl* with default settings. For gene ontology analysis, gene lists were analyzed for pathway and process enrichment using KEGG Pathways, GO Biological Processes, Reactome Gene Sets, Canonical Pathways, CORUM, and WikiPathways *via* Metascape^48^. All human genes were used as the enrichment background. Significant terms (*P*<0.01; minimum count = 3; enrichment factor >1.5 (enrichment factor is the ratio between the observed counts and the counts expected by chance)) were determined and clustered based on membership similarities (*P*-values were calculated based on the cumulative hypergeometric distribution, and *q*-values were calculated using the Benjamini-Hochberg procedure to account for multiple testing). Kappa scores were used as the similarity metric for hierarchical clustering of enriched terms, sub-trees with a similarity of>0.3 were considered a cluster, and the most statistically significant term within a cluster was chosen to represent the cluster.

### Statistical Analysis

Statistical analysis was performed and graphs were generated in GraphPad Prism 10.0.1 and R^49^. qRT-PCR data was analyzed using an unpaired, two-tailed *t*-test. *In vitro* proliferation data was analyzed using an unpaired, two-tailed *t*-test or one-way ANOVA with Tukey’s *post hoc* correction. *In vivo* tumor latencies were analyzed using one-way ANOVA with Tukey’s *post hoc* correction.

## ACKNOWLEDGEMENTS

This work was supported by the UCSF Brain Tumor SPORE Career Development Program, the UCSF Program for Breakthrough Biomedical Research (which is partially funded by the Sandler Foundation), the Shurl and Kay Curci Foundation, and NIH R01 AG064363 (B.S). B.S. was a Kimmel Scholar of The Sidney Kimmel Foundation and holds the Suzanne Marie Haderle and Robert Vincent Haderle Endowed Chair at UCSF. C.S. was supported by a PRCRP Horizon Award from the Congressionally Directed Medical Research Programs, U.S. Department of Defense. S.C. was supported by a Sullivan Postdoctoral Fellowship. W.A.W acknowledges R01NS125668; R01CA159859; R01CA255369; R01NS106155 and P30CA082103. This study was further supported, in part, by the HDFCCC Laboratory for Cell Analysis Shared Resource Facility through NIH grant P30CA082103. Parts of Figure 1A and Figure 6A were created with content licensed from *BioRender*.

## AUTHOR CONTRIBUTIONS

CS and BS designed and planned the study; CS, SH, AC, SC, SW, and LW performed research; CS, BLG, JJP, PAN, WAW, and BS analyzed data; WAW and BS supervised the research; CS and BS wrote the manuscript and all authors commented on the manuscript.

## COMPETING INTERESTS

B.S. is a member of the Scientific Advisory Board of Herophilus but declares no competing financial interest with the work reported here.

## SUPPLEMENTARY FIGURES

**Figure S1.**
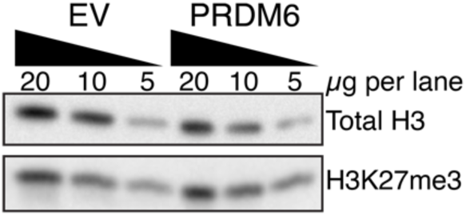
Global levels of H3K27me3 marks in PRDM6 and EV NES cells. The indicated amounts of acid-extracted histones were analyzed by immunoblotting with H3K27me3-specific antibodies. Blots were stripped and probed for total levels of histone H3.

**Figure S2.**
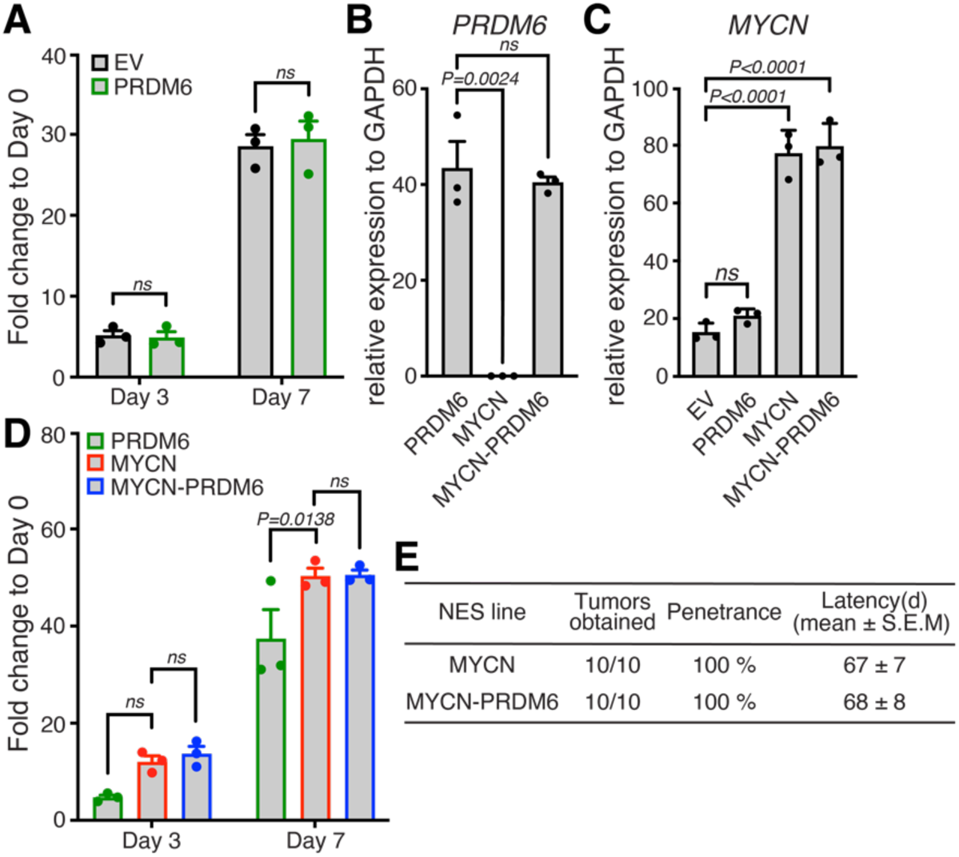
**(A)** *In vitro* proliferation assays of PRDM6 and EV NES cells on days three and seven. Error bars denote SEM. *P*-values were determined by one-way ANOVA with Tukey’s *post hoc* correction. **(B)** Quantification of *PRDM6* expression in PRDM6-, MYCN-, or MYCN-PRDM6 NES cells *via* qRT-PCR. Error bars denote SEM. *P*-values were determined by an unpaired, two-tailed *t*-test. **(C)** Quantification of *MYCN* expression, as described in (B). **(D)** *In vitro* proliferation of PRDM6, MYCN and MYCN-PRDM6 NES cells on days three and seven. Error bars denote SEM. *P*-values were determined by one-way ANOVA with Tukey’s *post hoc* correction. **(E)** Latency of tumor development after implantation of MYCN- or MYCN-PRDM6 NES cells into the cerebellum of mice. d, days.

**Figure S3.**
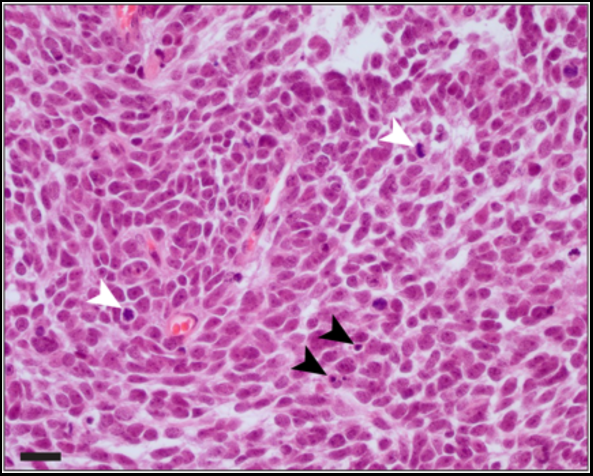
H&E staining of representative PRDM6 NES-derived tumor tissue. Mitotic figures (white arrowheads) and apoptotic bodies (black arrowheads) are highlighted, respectively. Scale bar, 20 μm.

## SUPPLEMENTARY TABLES

**Table S1.** Gene ontology analysis of regions with differential chromatin accessibility. (Excel file).

**Table S2.** Analysis of genomic regions with differential accessibility for known transcription factor binding motifs. (Excel file).

**Table S3.** Quantification of occupancy of transcription factor motifs in PRDM6 and EV NES cells. (Excel file).

**Table S4.** Genes overlapping or located within two kb of a PRDM6 binding site. (Excel file).

**Table S5.** *Molecular Signatures Database* (MSigDB) analysis of genes overlapping or located within two kb of a PRDM6 binding site. (Excel file).

